# Diversity and productivity of a natural grassland decline with the number of global change factors

**DOI:** 10.1101/2024.12.17.628835

**Authors:** Jianyong Wang, Yingxia Liu, Ayub M.O. Oduor, Mark van Kleunen, Yanjie Liu

**Author notes:** Corresponding author: Yanjie Liu, Email address.

## Abstract

Grasslands are highly diverse ecosystems providing important ecosystem services, but they currently face a variety of anthropogenic stressors simultaneously. Quantifying grassland responses to global change factors (GCFs) is crucial for developing effective strategies to mitigate the negative impacts of global change on grassland communities and to promote their resilience in the face of future environmental challenges. We conducted a field experiment in the Songnen grassland, northeastern China, to test the combined effects of 0, 1, 2, 4, 6, and 8 GCFs, including fungicide, herbicide, insecticide, antibiotic stress, heavy metal pollution, light pollution, microplastic pollution, nitrogen deposition, tillage disturbance, and increased precipitation. We found that within one year, the increasing number of GCFs negatively impacts on both the productivity and diversity of grassland communities. In comparison to exposure to a single GCF, exposure to 8 GCFs led to a reduction in productivity and species richness by 42.8% and 42.9%, respectively. Furthermore, these negative effects seem to be linked to the reduction of dominant species and the concurrent increase in neonative species. The results of hierarchical diversity-interaction modeling suggested that the adverse impacts of an increasing number of GCFs on community productivity and diversity are attributable to both the specific identities of GCFs involved and their unique pairwise interactions. The results suggest that grasslands may quickly loose stability and degrade more rapidly in response to multiple co-occurring GCFs. Greater efforts should be made to conserve the functions and services of grassland ecosystems by reducing the impacts of human activities.

## Introduction

Grasslands, encompassing a diverse range of ecosystems such as open grassland, grassy shrublands and savannas, cover a vast area of 52.5 million km², representing approximately 40.5% of the Earth’s land surface (excluding Greenland and Antarctica) (Bardgett et al. 2021). Grasslands are crucial for preserving biodiversity, hosting numerous iconic and endemic species, and offering a multitude of services to humans. These services include food production, water regulation, carbon sequestration for climate regulation, pollination, and numerous cultural services (Bardgett et al. 2021; Bai & Cotrufo 2022; Petermann & Buzhdygan 2021). In fact, around 1.5 billion people worldwide rely on grasslands for their livelihoods (Sun et al., 2022). Despite their vital importance as global reservoirs of biodiversity and providers of essential ecosystem services, grasslands are experiencing widespread and accelerating degradation across the globe due to ongoing and expanding human-caused global environmental changes (Bardgett et al., 2021).

Responses of grassland ecosystems to individual global change factors (GCFs) have been widely studied, providing valuable insights into how these ecosystems react to specific environmental changes. For instance, nitrogen enrichment, due to atmospheric nitrogen deposition and agricultural activities, has been shown to reduce plant species richness in grasslands by favoring fast-growing, competitive species over slower-growing, stress-tolerant ones (Suding et al., 2005). Similarly, changes in rainfall patterns, such as increased drought frequency or intensity, can alter competitive dynamics, leading to shifts in community composition (Wilcox et al., 2017). Even management practices like mowing can influence plant community structure by removing biomass and creating opportunities for less competitive species (Socher et al., 2012). However, the challenge lies in the fact that natural ecosystems rarely experience single isolated GCFs. Multiple GCFs, such as nitrogen enrichment, altered rainfall, and microplastic pollution, often act concurrently, creating a complex web of possible interactions (Rillig et al., 2019; Speißer et al., 2022; Xue et al., 2024).

The combined effects of multiple GCFs on grassland ecosystems can be additive, where the overall impact is simply the sum of the individual effects, synergistic, where the combined impact is greater than the sum of individual effects, or antagonistic, where the combined impact is less than the sum (Côté et al., 2016). Complex interactions can lead to unpredictable "ecological surprises", where the observed responses of grassland communities to multiple GCFs deviate significantly from expectations based on single-factor studies (Darling & Côté 2008). For example, the combined effects of nitrogen enrichment and altered rainfall patterns can cause more dramatic shifts in grassland community composition than would be predicted by considering each factor independently (Zavaleta et al., 2003). Such synergistic global change effects pose a substantial threat to biodiversity and ecosystem functioning (Oliver et al., 2015; van der Plas, 2019; Xu et al., 2024). Despite extensive knowledge on the responses of grasslands to single GCFs, our understanding of how multiple simultaneously acting factors affect these ecosystems remains limited (Zandalinas et al., 2021; Speißer et al., 2022). This knowledge gap is particularly concerning given the increasing prevalence of multiple, simultaneously acting GCFs (Steffen et al., 2015).

Multiple GCFs acting simultaneously can influence grassland communities through both direct and indirect pathways. Directly, the number of co-occurring GCFs can alter niche availability, either by shifting existing niches or creating new ones (Komatsu et al., 2019). This can lead to changes in plant community composition as species respond differently to the altered environmental conditions. Additionally, GCFs can directly impact plant growth, reproduction, and interactions with other species (Zandalinas et al., 2021; Zhou and Wang 2023). For instance, the presence of multiple stressors may weaken competition or promote facilitation among species, potentially increasing stress tolerance and influencing community dynamics. Furthermore, increasing numbers of GCFs can shift ecological tipping points, leading to abrupt transitions in ecosystem states (Suding & Hobbs 2009). Possibly, even low intensities of additional GCFs could already due to interactions with the other GCFs trigger a rapid ecosystem collapse (Zandalinas et al., 2021). Indirectly, the number of GCFs can alter top-down and bottom-up effects, impacting plant communities through trophic interactions and soil legacies. For example, GCFs that affect herbivory (e.g., insecticide, and light pollution) can influence plant communities via top-down effects by modifying herbivore behavior and abundance (Xu et al., 2024; Liu & Heinen 2024). Similarly, changes in the number of GCFs can alter interactions between plants and soil biota, thereby influencing plant communities (Xue et al., 2024). These altered plant-soil biota interactions can lead to reduced microbial diversity and shifts towards more generalist and stress-tolerant plant species (Rillig et al., 2019; Xu et al., 2024). While these mechanisms have been explored in various contexts, the impacts of multiple GCFs on natural grasslands remain largely unknown, highlighting the need for further research in this critical area.

Currently, studies exploring the responses of natural grassland communities to an increasing number of co-occurring GCFs are limited (Komatsu et al., 2019). Such studies, however, are crucial for developing effective strategies to conserve and manage ecosystems that are increasingly subjected to multiple environmental stressors (Rillig et al., 2019; Bowler et al., 2020). In particular, by conducting field experiments, researchers can gain valuable insights into the complex interactions between GCFs and their impacts on plant communities in real-world contexts. Therefore, we conducted a field experiment in a natural grassland to investigate how plant communities respond to an increasing number of simultaneously acting GCFs. We selected 10 GCFs that either already frequently occur simultaneously in grasslands or are predicted to become more common in the future: exposure to fungicide, herbicide, insecticide and antibiotics, heavy metal pollution, light pollution, microplastic pollution, nitrogen deposition, disturbance by tillage, and increased precipitation (Bardgett et al., 2021). We exposed natural grassland communities to one, two, four, six, or eight GCFs chosen randomly from this pool, alongside a control group with no GCFs. We addressed the question: Does the increasing number of GCFs affect the productivity, composition, and diversity of natural grassland communities?

## Results

### Effects of GCFs on productivity, total coverage, and plant height within the grassland community

The individual GCFs significantly impacted the productivity, total coverage, and plant height within the grassland community, but the direction and magnitude of these effects varied considerably among the GCFs (Fig. 1 & Table 1). Compared to the control, herbicide application and soil disturbance from tillage significantly decreased the mean community above-ground biomass per plot (Fig. 1a). Conversely, increased nitrogen input and precipitation significantly increased the mean community above-ground biomass (Fig. 1a). Moreover, herbicide application and soil disturbance from tillage significantly decreased, and fungicide application marginally decreased the community coverage per plot (Fig. 1b). Soil disturbance from tillage significantly increased mean plant height within the communities (Fig. 1c). As the number of simultaneously acting GCFs increased, we observed a significant decrease in both community productivity and total coverage (Fig. 1a, b). Conversely, plant height increased significantly with the number of GCFs (Fig. 1c).

**Fig 1.**
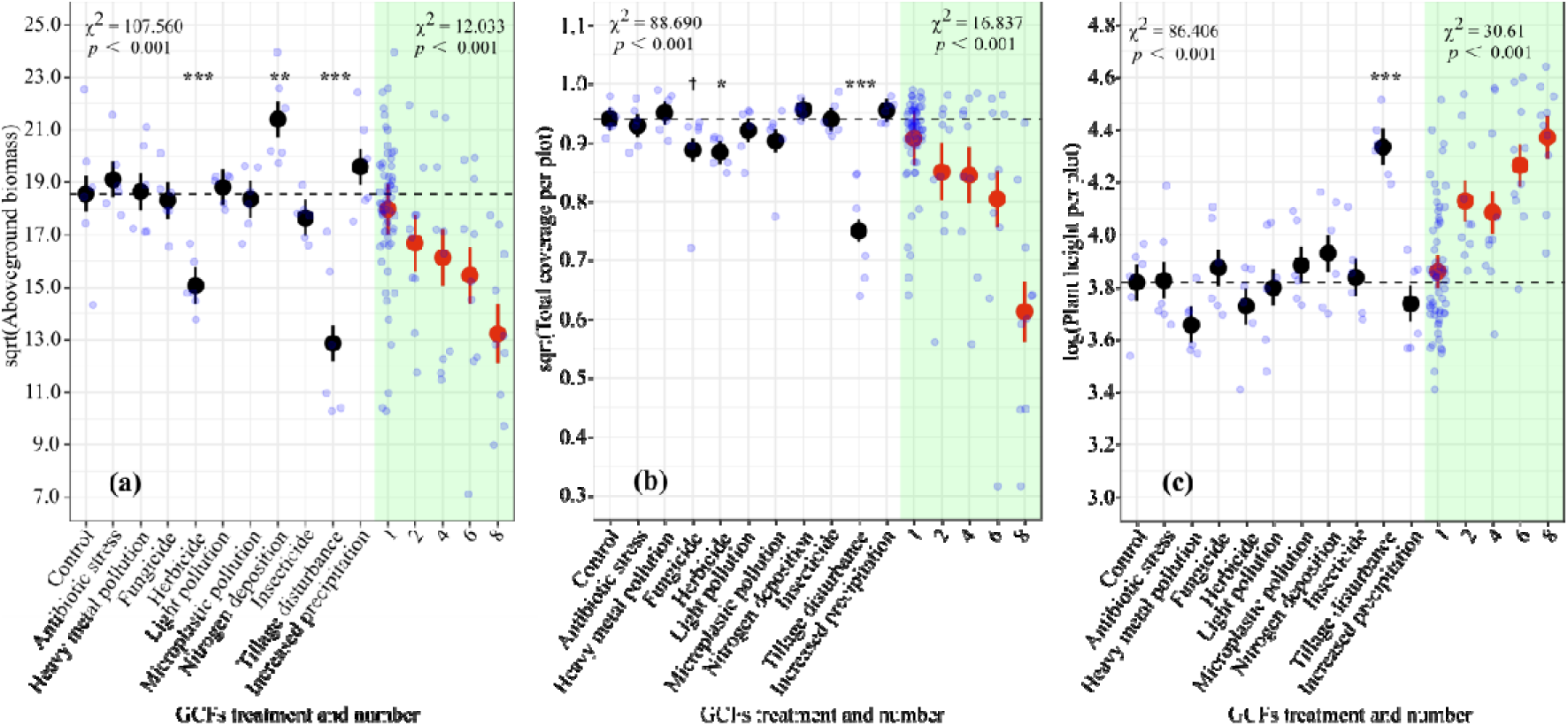
Effects of single and multiple global change factors (GCFs) on (a) community productivity, (b) vegetation coverage, and (c) community-weighted mean plant height in the Songnen grassland in northeastern China. Black circles represent responses to individual GCF treatments, while red circles represent responses to varying numbers of simultaneously acting GCFs (1, 2, 4, 6, and 8). The red circles marked with value “1” represent the average values across all individual GCF applications. Significant deviations of the individual GCF treatments from the control are indicated as follows: † (0.05 < *P* < 0.1), * (*P* < 0.05), ** (*P* < 0.01), and *** (*P* < 0.001). Error bars depict standard errors, and light blue circles show the raw data distribution. Horizontal dashed lines represent the mean values of the control group (no GCF treatment).

**Table 1.** Results of statistical tests that examined how the identity and interactions of global change factors (GCFs) influence various aspects of grassland plant communities, including productivity, coverage, plant height, diversity, and evenness.

| Model<br>(reference model number) | Above-ground biomass per plot |  |  |  |  | Total coverage per plot |  |  |  | Average plant height per plot |  |  |  |
| --- | --- | --- | --- | --- | --- | --- | --- | --- | --- | --- | --- | --- | --- |
| | DF | AIC | LogLik | $\chi^2$ | *P* | AIC | LogLik | $\chi^2$ | *P* | AIC | LogLik | $\chi^2$ | *P* |
| 1. Null | - | 104.55 | -49.28 | - | - | 84.93 | -39.46 | - | - | 9.64 | -1.82 | - | - |
| 2. Identity of factors (1) | 9 | 61.45 | -18.72 | 61.11<br>(+) | **<0.001** | 82.08 | -29.04 | 20.85<br>(+) | **0.013** | -26.42 | 25.21 | 54.06<br>(+) | **<0.001** |
| 3. Separate-pairwise factor interactions (4) | 37 | -15.03 | 57.52 | 152.40<br>(+) | **<0.001** | -96.21 | 98.10 | 252.72<br>(+) | **<0.001** | -33.58 | 66.79 | 78.48<br>(+) | **<0.001** |
| 4. Average factor interaction (2) | 1 | 63.37 | -18.69 | 0.08 | 0.781 | 82.52 | -28.26 | 1.56 | 0.212 | -29.10 | 27.55 | 4.69<br>(+) | **0.030** |
| 5. Additive factor-specific interaction contributions (3) | 28 | 47.64 | -1.82 | 118.68<br>(-) | **<0.001** | 67.60 | -11.80 | 219.8<br>(-) | **<0.001** | -35.23 | 39.61 | 54.36<br>(+) | **0.002** |

*Table 1 continued*
| Model<br>(reference model number) | Species number |  |  |  |  | Shannon-Wiener's diversity index |  |  |  | Simpson's diversity index |  |  |  |
| --- | --- | --- | --- | --- | --- | --- | --- | --- | --- | --- | --- | --- | --- |
| | DF | AIC | LogLik | $\chi^2$ | *p* | AIC | LogLik | $\chi^2$ | *p* | AIC | LogLik | $\chi^2$ | *p* |
| 1. Null | - | 74.58 | -34.29 | - | - | 18.27 | -6.13 | - | - | -94.66 | 50.33 | - | - |
| 2. Identity of factors (1) | 9 | 76.38 | -26.19 | 16.21 | 0.063 | 4.62 | 9.69 | 31.65<br>(+) | **<0.001** | -109.03 | 66.51 | 32.37<br>(+) | **<0.001** |
| 3. Separate-pairwise factor interactions (4) | 37 | 27.41 | 36.29 | 124.83<br>(+) | **<0.001** | -45.04 | 72.52 | 125.35<br>(+) | **<0.001** | -156.55 | 128.28 | 123.51<br>(+) | **<0.001** |
| 4. Average factor interaction (2) | 1 | 78.24 | -26.12 | 0.13 | 0.714 | 6.31 | 9.84 | 0.31 | 0.580 | -107.04 | 66.52 | 0.01 | 0.918 |
| 5. Additive factor-specific interaction contributions (3) | 28 | 60.42 | -8.21 | 89.01<br>(-) | **<0.001** | -10.65 | 27.33 | 90.38<br>(-) | **<0.001** | -132.40 | 88.20 | 80.15<br>(-) | **<0.001** |

*Table 1 continued*
| Model<br>(reference model number) | Pielou's evenness index |  |  |  |  |
| --- | --- | --- | --- | --- | --- |
| | DF | AIC | LogLik | $\chi^2$ | *p* |
| 1. Null | - | -88.60 | 47.30 | - | - |
| 2. Identity of factors (1) | 9 | -83.62 | 53.81 | 13.02 | 0.162 |
| 3. Separate-pairwise factor interactions (4) | 37 | -108.17 | 104.08 | 100.53 (+) | **<0.001** |
| 4. Average factor interaction (2) | 1 | -81.63 | 53.82 | 0.01 | 0.923 |
| 5. Additive factor-specific interaction contributions (3) | 28 | -87.90 | 65.95 | 76.27 (-) | **<0.001** |

Notes: Results are based on the hierarchical diversity-interaction comparisons between the model of interest and a reference model (indicated by the numbers in brackets) using log-likelihood ratio tests ( $\chi^2$ statistics). The numbers and names of the models correspond to those in Fig. S2. For each model, we show its AIC and log-likelihood (logLik). To answer the four questions shown in Fig. S2, we made four model comparisons based on log-likelihood ratio tests. The number of the reference model is shown in brackets after the model of interest. For the model comparisons, we provide the degrees of freedom (DF), the $\chi^2$ test statistic and the *p* value. The *p* values < 0.05 are indicated in bold. The + and – in brackets after the $\chi^2$ value indicate whether the model was better or worse than its reference model.

The hierarchical diversity-interaction modeling framework for productivity and total coverage revealed that the models accounting for separate pairwise GCF interactions provided the best fit (Table 1). This suggests that the negative effects of increasing GCF number on community productivity and coverage are attributable to both the specific identities of the GCFs involved and their unique pairwise interactions. In contrast, for the mean plant community height, the model incorporating additive factor-specific interaction contributions showed the best fit (Table 1). This suggests that the positive effect of increasing the number of global change factors (GCFs) on plant height was driven by specific interactions between these factors.

### Effects of GCFs on plant community composition and diversity

Single-factor treatments significantly influenced species richness and diversity (Shannon-Wiener and Simpson indices) within the grassland community (Fig. 2), but had no effect on evenness. Soil disturbance from tillage significantly decreased species richness, and the Shannon-Wiener and Simpson indices (Fig. 2a-c). Moreover, light pollution significantly reduced the Shannon-Wiener and Simpson indices (Fig. 2b, c). Fungicide application tended to decrease species richness (Fig. 2a). Mirroring the trends observed in community productivity, we found significant directional changes linked to the number of GCFs: a reduction in species richness, evenness, Shannon-Wiener index, and Simpson index (Fig. 2). The identity and number of GCFs both had significant effects on plant community composition (Fig. 3a, b).

**Fig 2.**
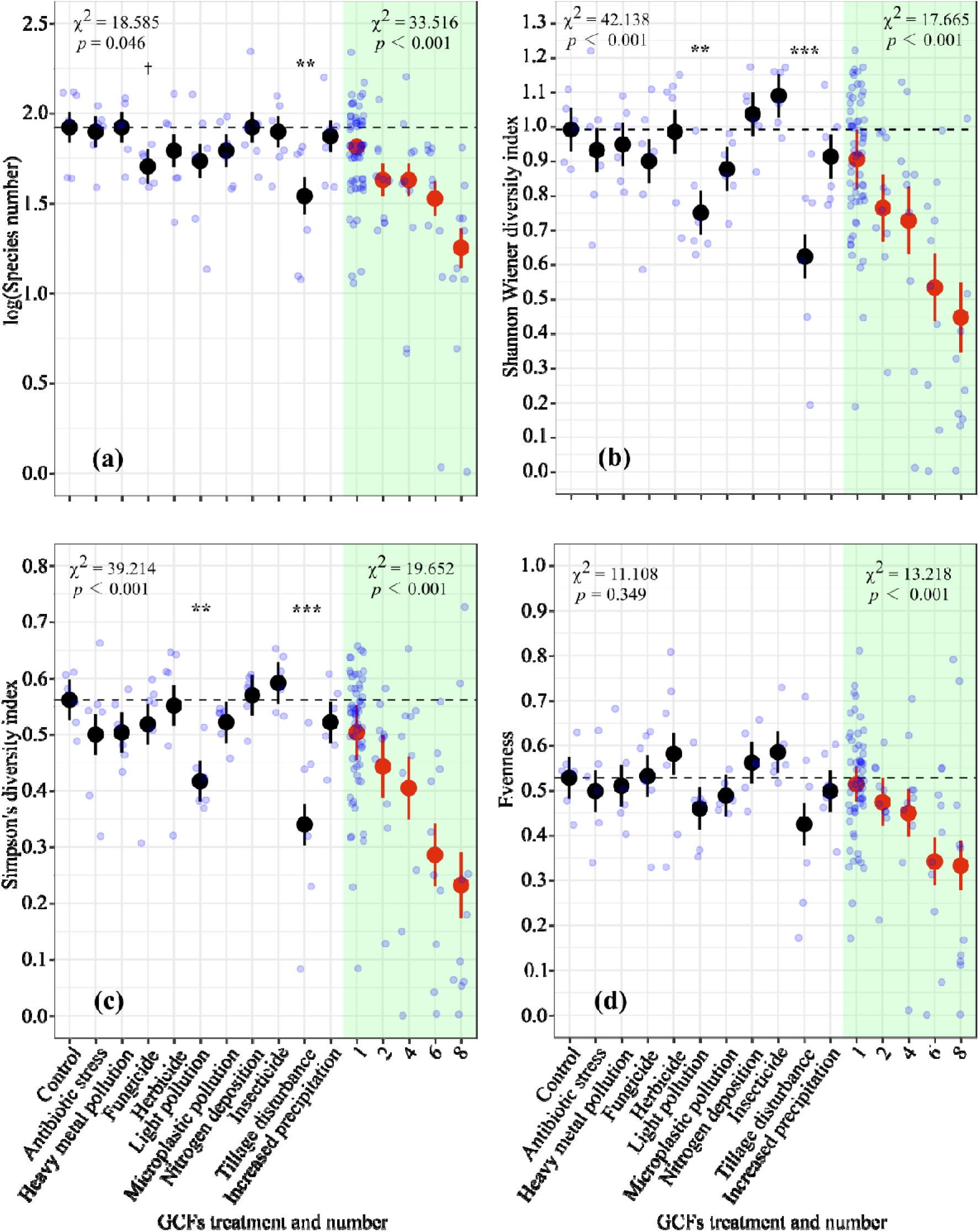
Effects of single and multiple global change factors (GCFs) on (a) species richness, (b) Shannon-Wiener diversity index, (c) Simpson’s diversity index, and (d) Evenness in the Songnen grassland in northeastern China. Black circles represent responses to individual GCF treatments, while red circles represent responses to varying numbers of simultaneously acting GCFs (1, 2, 4, 6, and 8). The red circles marked with value “1” represent the average values across all individual GCF applications. Significant deviations of the individual GCF treatments from the control are indicated as follows: † (0.05 < *p* < 0.1), * (*p* < 0.05), ** (*p* < 0.01), and *** (*p* < 0.001). Error bars depict standard errors, and light blue circles show the raw data distribution. Horizontal dashed lines represent the mean values of the control group (no GCF treatment).

**Fig 3.**
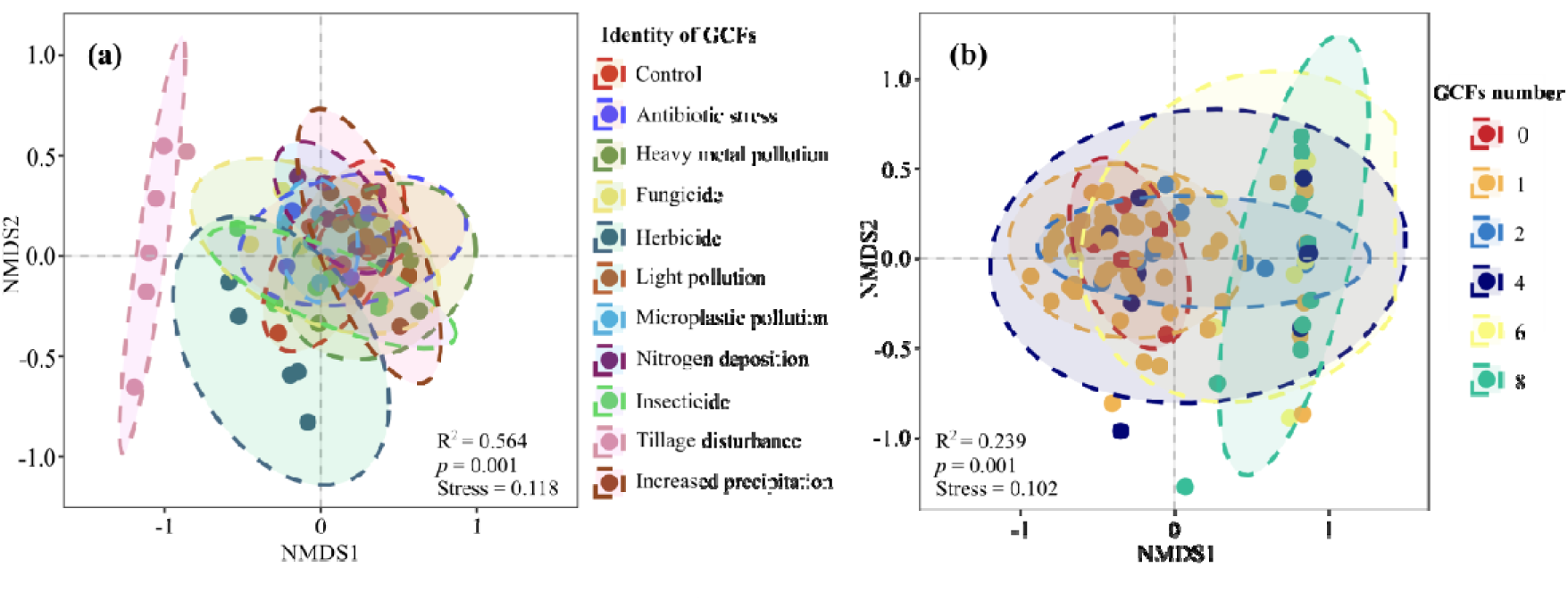
Nonmetric multidimensional scaling (NMDS) plots showing the grassland community composition under (a) different single global change factors (GCFs), and (b) under treatment combinations involving one, two, four, six, and eight GCFs and a control without any GCF.

Applying the hierarchical diversity-interaction modeling framework, we found the best-fitting model for species richness, diversity, and evenness to be the one incorporating separate pairwise GCF interactions (Table 1). This suggests that the negative impact of increasing GCF number on these variables stems from both the identities of the GCFs themselves and their strong, specific pairwise interactions.

## Discussion

Our field experiment in a natural grassland showed that most of the 10 GCFs applied individually had weak or minimal impacts on the productivity, species composition, and diversity of the grassland community. Additionally, the effects of single GCFs varied depending on the specific community variables examined (Figs. 1, 2). However, despite the weak individual effects of most GCFs, our study showed that an increase in the number of GCFs led to more pronounced directional changes in the grassland-community attributes. Consistent with previous mesocosm studies (Rillig et al., 2019; Speißer et al., 2022; Yang et al., 2022), our findings thus indicate that the presence of multiple GCFs acting in concert can lead to more severe consequences for grassland communities compared to the additive effects of individual factors. The synergistic interactions among GCFs potentially lead to more rapid and extensive degradation of grassland productivity and diversity, and changes in species composition. These findings align with the growing body of evidence that underscore the importance of considering the joint effects of multiple GCFs when assessing the vulnerability of ecosystems to global change (Orr et al., 2020).

Previous research has demonstrated that an increasing number of simultaneously acting GCFs can negatively affect soil properties, soil microbial diversity, and the growth, survival and diversity of plants (Rillig et al., 2019; Zandalinas et al., 2021; Speißer et al., 2021; Yang et al., 2022; Fu et al. 2024). A more recent study also found that increasing numbers of GCFs can alter plant-soil feedbacks by altering the fungal legacies left by plants in the soil (Xue et al., 2024). These studies underscore the critical role of multiple GCFs in influencing plants and soils and regulating ecological processes such as plant-microbe interactions, which in turn can impact community structure and functions. Our study builds upon this existing knowledge from artificial microcosm and mesocosm studies by providing the perspective from a natural grassland community. We demonstrate that the increasing number of co-occurring GCFs does not only affect individual plants and soil components but also leads to a significant decrease in the productivity and diversity of entire grassland communities.

Our study’s findings differ from those of a previous study by Speißer et al. (2022), which found that an increasing number of GCFs led to greater productivity in a grassland mesocosm. This was primarily driven by a strong positive influence of nitrogen enrichment in that study, which was, due to the limited pool of GCFs in that study, more likely to be included in the high GCF-number treatments. Moreover, in contrast to Speißer et al. (2022), our study was conducted in a natural grassland where complex conditions may weaken the concurrent effects of N addition with other GCFs on plant communities. Many previous studies have been conducted under greenhouse conditions or in experimental mesocosms (Speißer et al., 2022; Yang et al., 2022), where the experimental conditions do not fully mirror the natural habitats of the plants and may not accurately represent the complex interactions and feedbacks that occur in real-world ecosystems (De Boeck et al., 2015; Leuzinger et al., 2011). To address the gap in knowledge on real-world impacts of multiple global change factors (GCFs), long-term, field-based studies in natural habitats are crucial. This research will enhance our understanding of the cumulative effects of GCFs on biodiversity, ecosystem functioning, and services, while also revealing potential tipping points, regime shifts, and the overall resilience of ecosystems to global change (Blowes et al. 2019; Ratajczak et al. 2018).

The strong effects of the number of GCFs on the plant community in our study may have arisen for several reasons. First, changes in resource availability may affect niche dimensionality, with large consequences for plant communities. For example, light pollution and N enrichment may decrease the heterogeneity in the availability of limiting resources, thereby decreasing the amount of niche space for coexistence without necessarily reducing the number of limiting resources (Harpole & Tilman 2007). Combinations of GCFs that synergistically enhance resource availability can reduce niche dimensionality, while combinations that decrease resource availability can increase it. Second, an increasing number of GCFs increases the likelihood of including at least one GCF with strong effects (i.e., a sampling effect). Light pollution, for instance, which had strong negative effects on several of the diversity metrics, can alter plant communities via effects on herbivores and via increased resource availability. Although studying every possible interaction among the many different global change factors is unfeasible, our results are consistent with a recent meta-analysis reporting that plant communities were more likely to be altered when facing at least three global change factors acting simultaneously (Cheng et al., 2024). Third, co-acting GCFs may alter the individual effects of other factors, with joint net effects depending on whether and how these factors interact (interaction effect). In our study, hierarchical diversity-interaction models indicated that in addition to the effect of GCF identities (e.g., N enrichment or tillage), specific pairwise interactions between the global change factors also contributed to the observed negative effects of increasing numbers of GCFs on productivity and diversity of the grassland community (Table 1). This means that the effect of multiple GCFs on ecosystems might deviate considerably from predictions based on the effect of single GCFs.

Our results indicate that multiple, simultaneously applied GCFs differentially impacted *Leymus chinensis*, which is the dominant species in this grassland, and *Phragmites australis*. The latter is native to most of China, but has recently started to expand its range into the meadow steppe where our study site is (i.e. it is a neonative *sensu* Essl et al., 2021). While the biomass of the dominant species declined with increasing GCF numbers, the neonative species increased its biomass proportion. Consequently, the overall negative effect of multiple GCFs on community biomass appears to be driven by the decline of the dominant species and the simultaneous rise of the neonative species. Plant community composition varied significantly across different single GCF treatments, with tillage exerting the most pronounced impact (Fig. 3a). This disturbance favored the growth of *P. australis*, which is a much taller species than the others species in the grassland community. *Phragmites australis* is a widely distributed and highly polymorphic species that is primarily found along riverbanks and lake margins, where it plays a crucial role in maintaining wetland ecosystem diversity (An et al., 2012). However, it also thrives in arid environments, such as deserts and dry lands in western and northwestern China (An et al., 2012). The increased abundance of this tall invasive species following tillage likely explains the observed positive correlation between the number of GCFs and overall plant height at the community level.

In summary, this study is the first to test the joint effects of multiple co-occurring GCFs on a natural grassland community. We found that an increasing number of GCFs can reduce grassland productivity and biodiversity through both individual GCF effects and their specific interactions. Notably, these effects were observed within a single year, indicating that natural ecosystems may be more sensitive to GCFs than previously thought. This underscores the importance of conducting studies that investigate the impacts of co-occurring multiple GCFs on natural ecosystems, as revealed by recent research (Rillig et al., 2019; Speißer et al., 2022; Xu et al., 2024). To advance this research, future studies should prioritize long-term field experiments in natural habitats to capture the temporal dynamics of grassland responses to multiple GCFs.

## Materials and Methods

### Site description

We conducted the experiment in the Songnen grassland, located in Changling County (44° 45′N, 123°45′E), Jilin Province, in northeastern China. This region experiences a semiarid continental and monsoonal climate characterized by cold, dry winters and warm, moist summers. The average annual temperature ranges from 4.6 to 6.4 °C, while annual precipitation varies between 253.2 and 716.2 mm, with approximately 70% falling during the growing season from July to September (Wang 2023). Soil salinization and alkalization are the main limiting factors for vegetation development in this area. The major vegetation type is meadow steppe dominated by *Leymus chinensis* (Trin.) Tzvel.

### Experimental design and plot lay-outs

The chosen GCFs encompass a broad range of environmental stressors, including: antibiotic exposure, heavy metal pollution, fungicide accumulation, herbicide accumulation, insecticide accumulation, light pollution, microplastic pollution, N deposition, disturbance by tillage, and increased precipitation. Each GCF represents a distinct aspect of global environmental change: resource availability (nitrogen deposition, increased precipitation, light pollution), chemical pollutants (antibiotics, heavy metals, fungicides, herbicides, pesticides), a physically and potentially chemically acting agent (microplastics), and a physical disturbance (tillage) (Rillig et al., 2019; Speißer et al., 2022; Xue et al., 2024). We selected these GCFs due to their prevalence in the region of study and their distinct modes of action and potentially varied impacts on grassland ecosystems.

To investigate the effects of increasing numbers of co-occurring GCFs on grassland plant communities, we designed six levels of GCF diversity. These levels included zero (control with no GCF), one (a single GCF), two, four, six and eight GCFs (*sensu* Rillig et al., 2019; Speißer et al., 2021). We randomly assigned individual GCFs from our pool of 10 to these groups. The control and single GCF groups had six replicates each. For groups with two, four, six, or eight GCFs, we randomly created combinations, while ensuring that each GCF was evenly represented across the different factor-number levels, with 10 replicates each (Table S1). This experimental design parallels methods used in biodiversity research (e.g., Tilman et al., 1997), focusing on GCF diversity rather than the specific identity or combination of GCFs (Rillig et al., 2019). This design resulted in a total of 106 experimental plots (Fig. S1a), each measuring 2.8 m × 2.8 m. To prevent interference of the treatment applications between plots, a 2.0-m buffer zone separated each plot. In addition, to contain the GCFs within their designated plots, we enclosed each plot with iron sheets buried 40 cm deep and extending 20 cm aboveground (Fig. S1b).

### The implementation of GCFs

Between May 11 and June 25, 2023, we implemented the 10 GCFs in various combinations to the plots. For the antibiotic stress treatment, 3.5 mg of oxytetracycline per kg of soil was added to the top 20 cm of soil, resulting in a final concentration of 3.5 mg/kg. This dosage is within the range used in previous studies and reflects antibiotic exposure levels common in agriculture. We dissolved 9.42 g of oxytetracycline in 3 liters of water and distributed it evenly over the soil surface using a watering can. For the microplastic pollution treatment, we used polybutylene succinate (PBS) granules (3.0–4.0 mm in diameter) at a concentration of 0.1% (w/w, granules/dry soil). PBS is commonly found in packaging, medical supplies, agricultural mulching, and household items, making it relevant for studying microplastic effects on plants. In each designated plot, 2.60 kg of PBS was sprinkled evenly over the soil surface. For the heavy metal (cadmium) pollution treatment, CdCl_2_ was added to the top 20 cm of the soil. We dissolved 26.13 g of CdCl_2_ in 3 liters of water and evenly distributed it over the soil surface, achieving a concentration of 10.0 mg/kg. For pollution by fungicides, 0.9 g of Score® (10% difenoconazole) and 0.225 g of Ortiva® (250 g/L azoxystrobin) were dissolved in 0.5 liters of water and sprayed evenly onto the plants in each plot. For pollution by a herbicide, 0.235 ml of glyphosate (30%) was diluted in 0.5 liters of water and sprayed onto the plants. Insecticide application involved spraying 0.27 ml of Karate Zeon® (25 g/L lambda-cyhalothrin) diluted in 0.5 liters of water onto the plants. For the nitrogen deposition treatment, 169.7 g of slow-release urea (46.2% N) was evenly spread in each plot, corresponding to an input of 10.0 g N/m², similar to the annual atmospheric nitrogen deposition in northern China (Wen et al., 2020). For the light-pollution treatment, LED spotlights mounted on 120 cm high metal frames were used (Fig. S1c), providing an average light intensity of 30.0 lux at ground level from sunset to sunrise, which is within the range of light intensities found below light poles (Speißer et al. 2022). To achieve full inversion tillage, a small rotary tiller was used to plow the entire plot to a depth of 20 cm (Fig. S1d). For the increased precipitation treatment, 258.4 liters of water were added per plot, simulating a 30% increase above the average rainfall received between May and September over the past two decades in this region (Wang 2023). For each treatment, in which solutions were applied to the respective plots, an equivalent amount of water was applied to the control plots. A summary of the levels of the individual GCFs used in this experiment is provided in Supplementary Table S2.

### Measurements

We carried out sampling and measurements in all the 106 plots on October 3, 2023 (i.e. 4 months after the start of the GCF treatments). To assess above-ground vegetation biomass, we randomly placed a 50 cm × 200 cm quadrat in each plot and clipped all vegetation at ground level. Immediately after clipping, we identified each plant species present in the sample. We then randomly selected five harvested individuals of each species and measured their height. The collected plant material was placed in paper bags, flash-dried at 105 °C in a forced-air oven for 2 hours, and then oven-dried at 65 °C for 72 hours until a constant weight was achieved.

We assessed plant diversity and composition in each of the 106 plots. Plot-level plant species richness (*S*) was calculated by counting the total number of species present in the sampled quadrat. We also calculated plot-level diversity using the Shannon-Wiener diversity index (H’) and Simpson’s diversity index (D), and evenness using Pielou’s evenness (J = H’/ln[S]), based on the total aboveground biomass of each species within the sampled quadrat. These calculations were done using the diversity function in the "vegan" package (Oksanen et al., 2022). We also estimated total vegetation coverage in each plot. As plant height is a key indicator of light utilization strategy and competitive ability, we calculated the community-weighted mean (CWM) of plant height using the following equation:

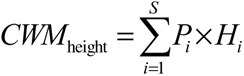

Where S is the species richness, H_i_ is the average height of 5 individuals of species i, and Pi represents the relative abundance of species i.

### Statistical analyses

All statistical analyses were done in R 4.2.2 (R Core Team 2022). To examine the combined effects of GCFs on community productivity, we fitted linear mixed-effects models using the *lme* function in the “nlme” package (Pinheiro et al., 2021). The above-ground biomass per plot was the response variable. To improve normality of the residuals, the above-ground biomass was square-root transformed. Additionally, we assessed the effects of individual GCFs on total biomass using linear mixed-effects models, incorporating only the control and single-GCF treatment data, with GCF identity as a fixed effect. To analyze the effects of single GCFs and increasing GCF numbers on plant community composition, diversity, and evenness, we also used linear mixed-effects models. The significance of fixed effects in all models was evaluated using Type II analysis of variance (ANOVA) with the *Anova* function from the "car" package.

A significant effect observed in relation to the number of GCFs could be due to the increased likelihood of including a dominant GCF when more factors are present, a phenomenon known as the “sampling effect”. However, this effect could also stem from interactions among multiple GCFs, where these interactions might either be specific to certain GCFs or represent a more generalized effect. Since our experiment did not encompass all possible combinations of GCFs, we could not directly test the impact of each potential multi-way GCF interaction. Therefore, to determine whether GCF identities or specific/general interactions underlie the significant GCF-number effects observed, we employed the hierarchical diversity-interaction modeling framework developed by Kirwan et al. (2009), specifically adapted to GCF identities and interactions. For each response variable with a significant GCF-number effect, we ran five hierarchical models, each with different assumptions regarding the contributions of single GCFs and their interactions. We then compared these models using likelihood-ratio tests, and the best model was determined using AIC (Fig. S2 & Table 1). This approach allowed us to discern the relative importance of GCF identity versus interactions in driving the observed responses of grassland communities to increasing GCF numbers.

We also performed PERMANOVA, using the *adonis2* function in the *vegan* package, to test whether plant community composition could be explained by the number and the identity of GCFs; significance was based on a permutation test (999 permutations). Differences in plant community composition among the GCF treatments were visualized using non-metric multidimensional scaling (NMDS) plots based on the Bray-Curtis dissimilarity matrix (Schroeder & Jenkins 2018).

## Acknowledgements

Yanjie Liu acknowledges funding from Innovation Team Project of Northeast Institute of Geography and Agroecology, Chinese Academy of Sciences (2022CXTD01). Ayub M. O. Oduor acknowledges funding from the Chinese Academy of Sciences–President’s International Fellowship Initiative (CAS-PIFI) (2021VBB0004). Jianyong Wang acknowledges funding from the National Natural Science Foundation of China (32271585).

## Supporting information

**Figure S1.**
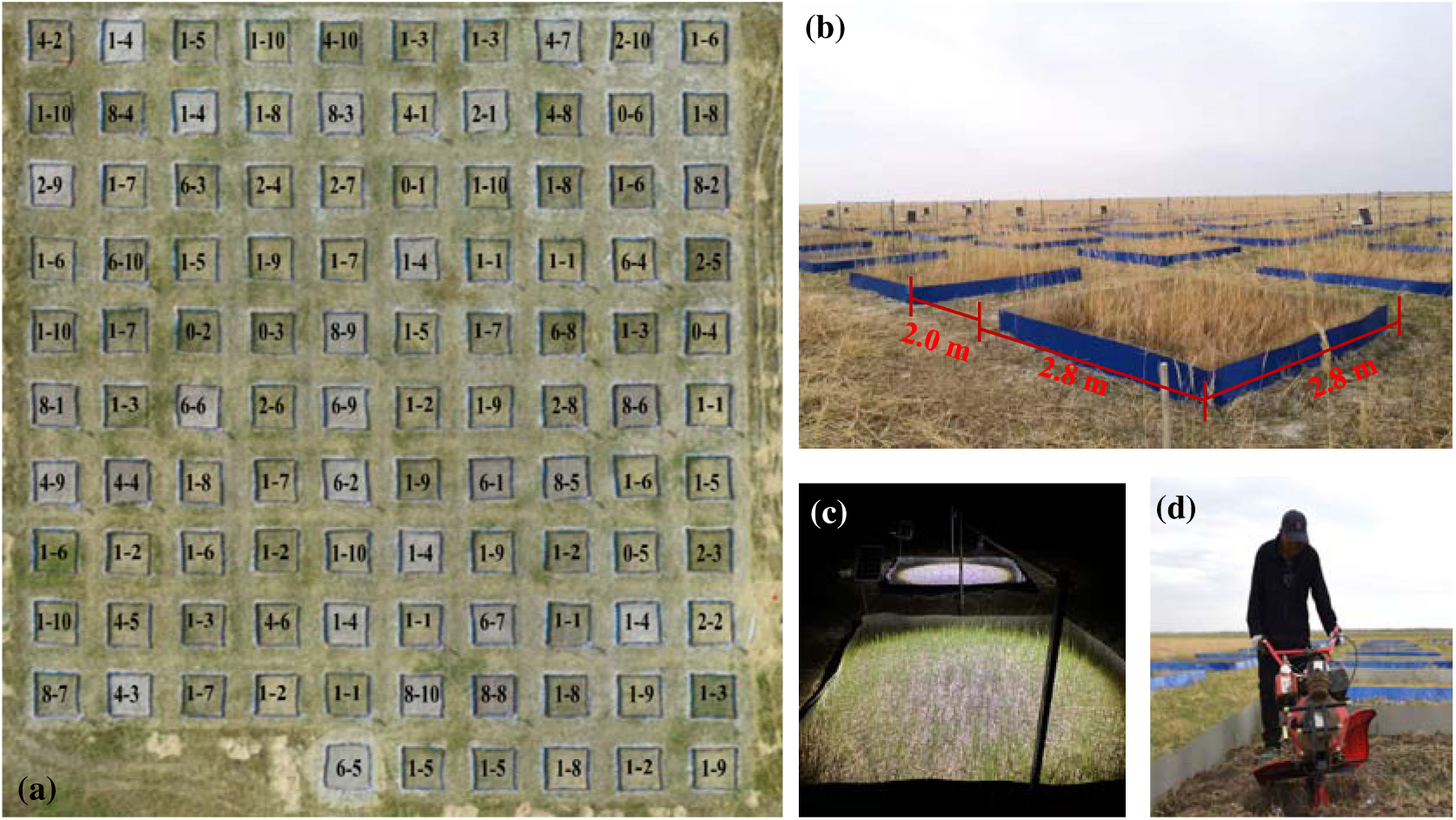
Overview of the experimental plot layout. (a) Distributions of the 106 plots in this study, with their treatment-codes corresponds to those in Table S1. (b) Each plot measures 2.8 m × 2.8 m, with a 2.0 m buffer between plots. (c) Light-pollution treatment. (d) Tillage disturbance treatment.

**Figure S2.**
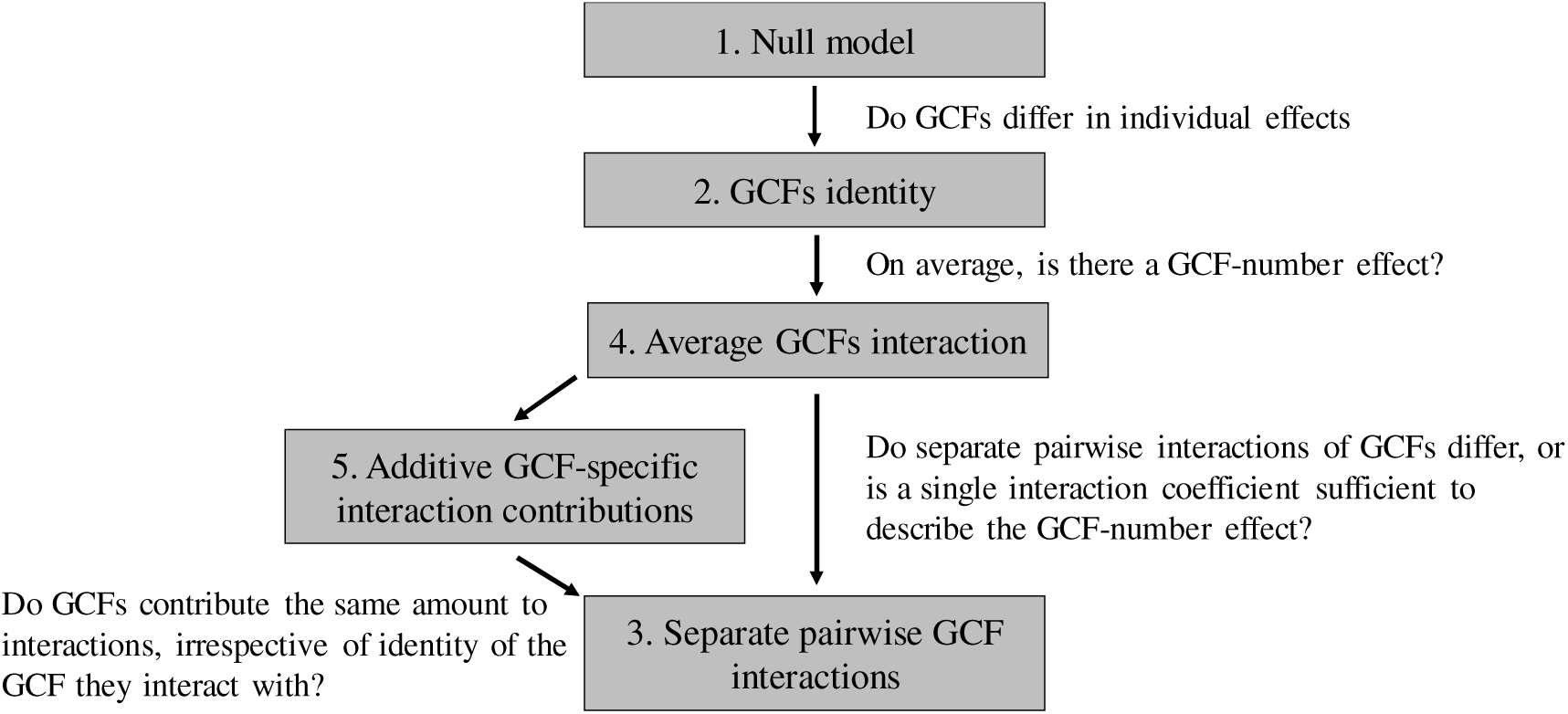
Hierarchical framework for analyzing individual and interactive effects of global change factors (GCFs). The framework was adapted from Kirwan et al. (2009). The null model assumes equal contributions of all GCFs and no interactions between them, while subsequent models assume more complex effects of the single GCFs and their interactions determine the GCF-number effects. The specific questions that can be answered by comparing specific models are depicted next to the arrows connecting them.

**Figure S3.**
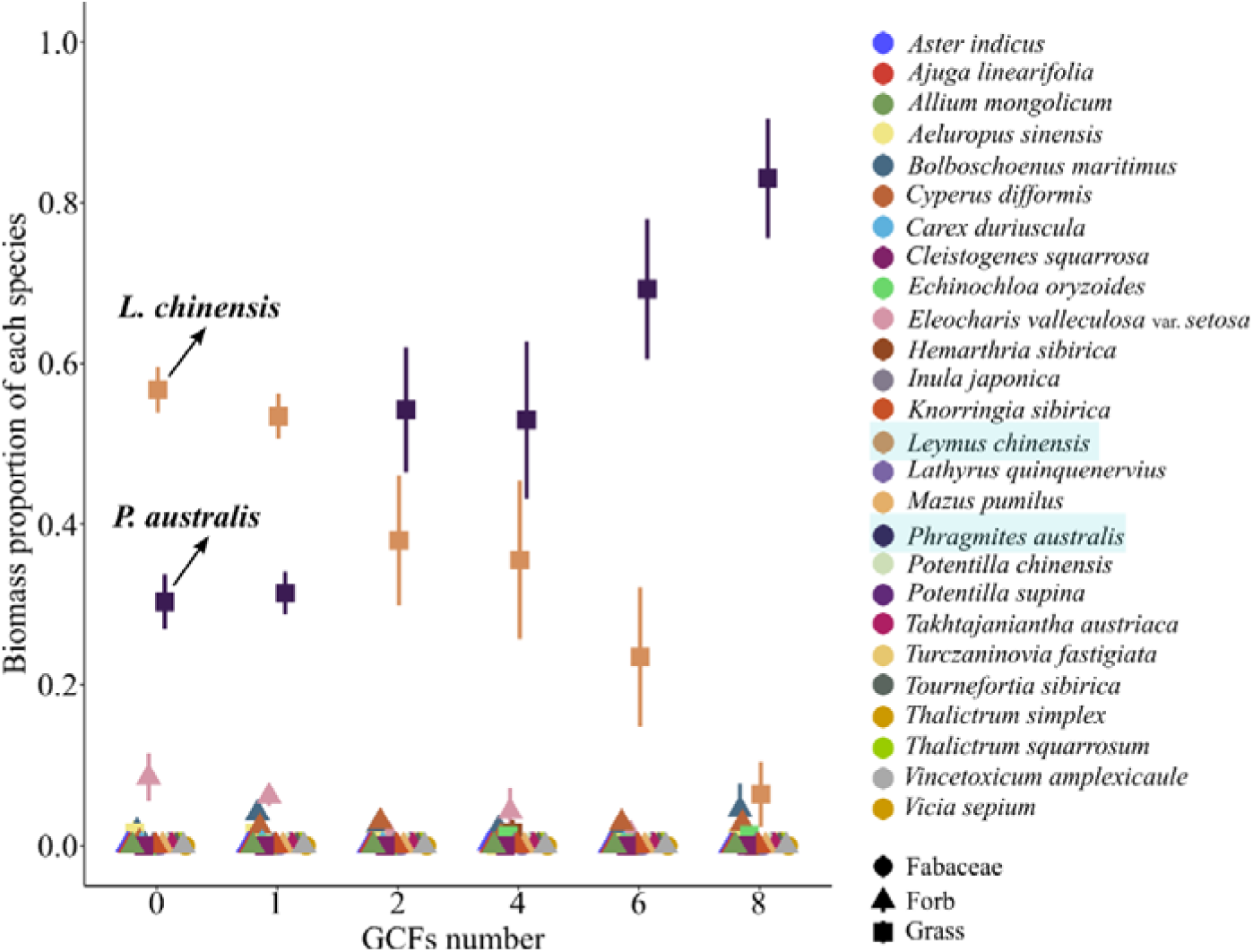
The effects of the number of simultaneously acting global change factors (GCFs) on the mean biomass proportions of different plant species within communities were investigated. The biomass proportions of each species were analyzed in relation to the number of co-occurring GCFs. Mean values are represented by symbols, with symbol shapes indicating the functional group of the species and error bars representing standard errors. The sample sizes (n) for each GCF-number level were as follows: n = 6 for zero GCFs, n = 60 for one GCF, and n = 10 for two, four, six and eight GCFs. The two species highlighted in blue color in the legend are the locally dominant plant species, *Leymus chinensis*, and the neonative species, *Phragmites australis*, respectively.

**Table S1.**
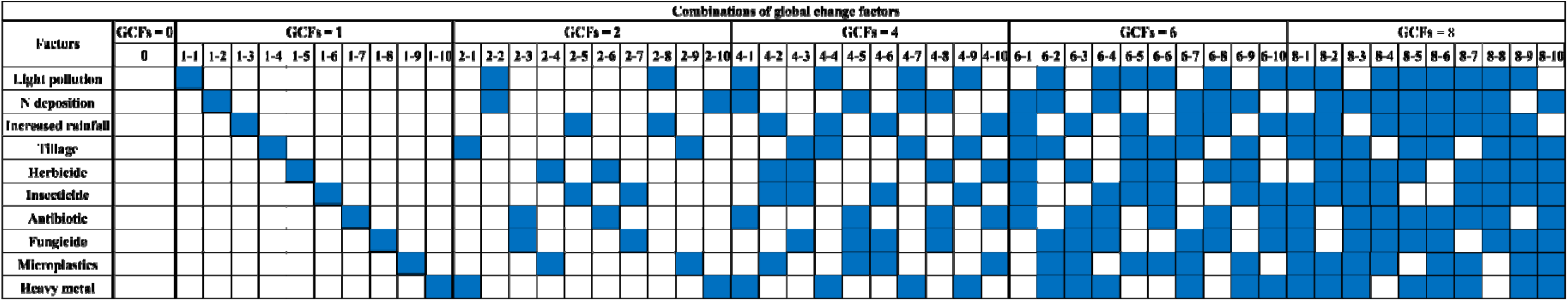
Combinations of global change factors (GCFs) applied at various levels in this experiment. Treatments involving zero or one GCF were replicated six times (n=6), while treatments involving two, four, six, or eight GCFs were replicated ten times (n=10). Blue shading indicates the presence of a specific GCF in the respective GCF diversity level. The number from 1 to 8–10 represents the treatment-code.

**Table S2.**
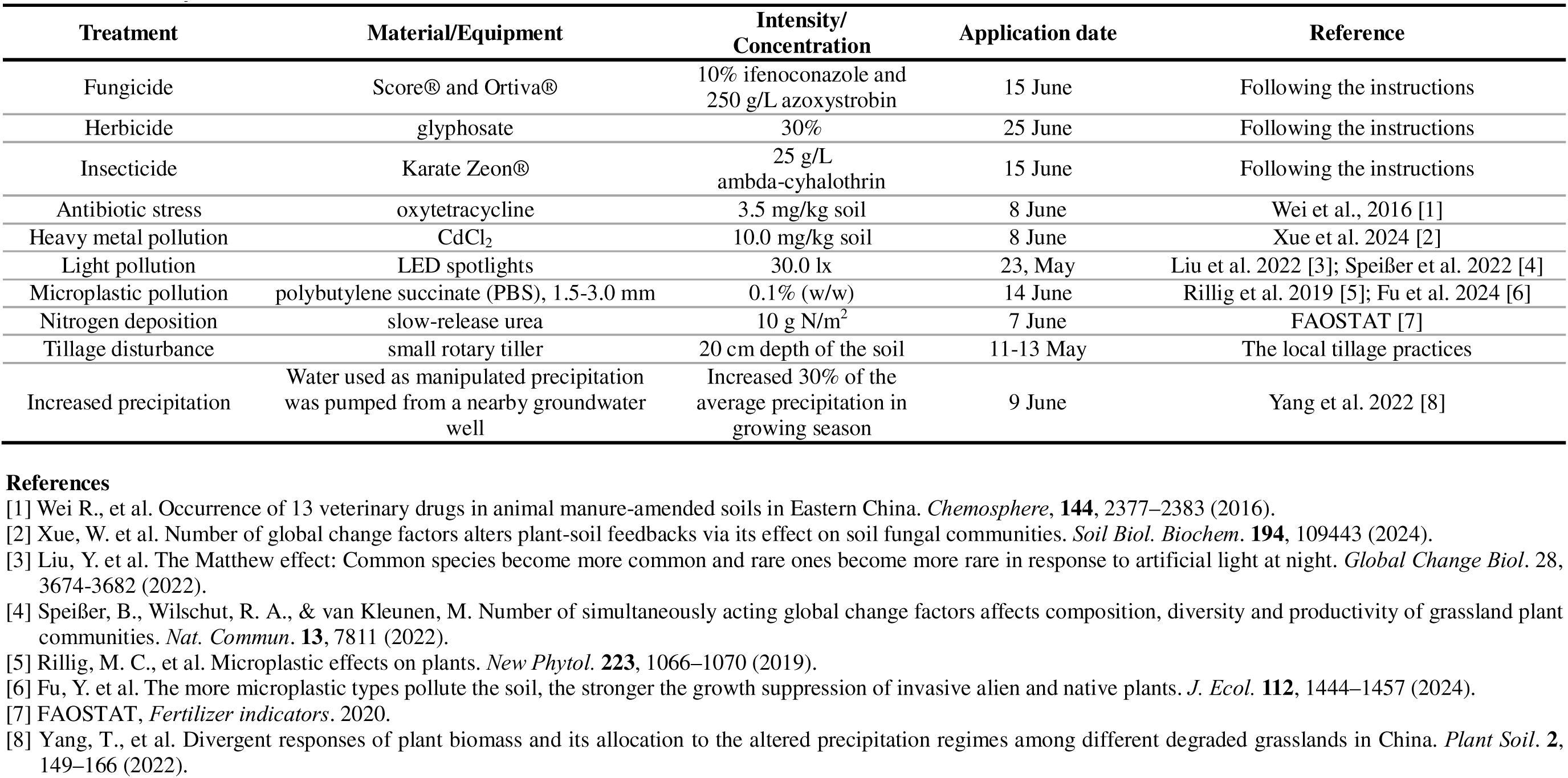
Single GCF treatment implementation. Summary of the materials or equipment used to implement the single GCF treatments, and the intensity of concentration of each GCF.

